# Contrasting stripes are a widespread feature of group living in birds, mammals and fishes

**DOI:** 10.1101/2020.04.20.050245

**Authors:** Juan J. Negro, Jorge Doña, M. del Carmen Blázquez, Airam Rodríguez, James E. Herbert-Read, M. de L. Brooke

**Author notes:** Corresponding author: Juan J. Negro.

## Abstract

Grouping is a widespread form of predator defense, with individuals in groups often performing evasive collective movements in response to predators’ attacks. Individuals in these groups use behavioral rules to coordinate their movements, with visual cues about neighbors’ positions and orientations informing movement decisions. Although the exact visual cues individuals use to coordinate their movements with neighbors have not yet been decoded, some studies have suggested that stripes, lines or other body patterns may act as conspicuous conveyors of movement information that could promote coordinated group movement, or promote dazzle camouflage, thereby confusing predators. We used phylogenetic logistic regressions to test whether the contrasting achromatic stripes present in four different taxa vulnerable to predation, including species within two orders of birds (Anseriformes and Charadriiformes), a suborder of Artiodactyla (the ruminants) and several orders of marine fish (predominantly Perciformes) were associated with group living. Contrasting patterns were significantly more prevalent in social species, and tended to be absent in solitary species or species less vulnerable to predation. We suggest that stripes taking the form of light-colored lines on dark backgrounds, or *vice vers*a, provide a widespread mechanism across taxa that serves either to inform conspecifics of neighbors’ directional movement, or to confuse predators, when moving in groups. Detection and processing of patterns and of motion in the visual channel is essentially colourblind. That diverse animal taxa with widely different vision systems (including di-, tri- and tetrachromats) appear to have converged on a similar use of achromatic patterns is therefore expected given signal-detection theory. This hypothesis would explain the convergent evolution of conspicuous achromatic patterns as an antipredator mechanism in numerous vertebrate species.

## Introduction

Group living characterizes the lives of many animals including flocking birds, schooling fish and ungulate herds. Grouping is often a defensive strategy to reduce the likelihood of individuals being attacked through detection, dilution, confusion, or self-herd effects (Hamilton 1971, Foster and Treherne 1981, Lima 1995, Jeschke and Tollrian 2007). Because isolated individuals are more likely to be targeted by predators (Ioannou 2012, Hogan et al. 2017, Cuthill 2019, Romenskyy et al. 2020), individuals in groups often move to maintain cohesion by escaping from predators in the same direction as other group members (Handegard et al. 2012, Herbert-Read et al. 2015). Many models aimed at explaining such coordinate movements assume animals rely primarily on visual information to infer neighbors’ movements and positions (Pita et al. 2016). These assumptions seem appropriate considering that birds are highly visually-oriented animals (Martin 2017, Camacho et al. 2019), and many fishes and mammals similarly rely on vision when interacting with conspecifics and their surroundings over intermediate distances (Guthrie and Muntz 1993).

Because vision appears essential for many species relying on coordinated and cohesive escape from predators, particular visual cues may be used by social animals to promote coordination. Numerous grouping species display conspicuous stripes on their bodies (Figure 1), with differences in design related to their particular body plans. In mammals and fishes, the majority of which have elongated bodies parallel to the substrate, stripes often run parallel to the longitudinal axis of the body. In birds, contrasting lines are often displayed in open wings. In all groups, stripes are typically achromatic: light-colored lines, often pure white, appear on an otherwise darker (i.e., melanized) plumage. Brooke (1998) found that flocking shorebirds tended to have white wing stripes, while solitary species did not, and suggested that these white marks could provide a conspicuous flash to other group members promoting cohesive take-off, or could aid in coordinated movement. In such species, a role of sexual selection as a driver of such body patterns may be dismissed, as these marks are shown year-round by both males and females in otherwise sexually dichromatic species that display both breeding and non-breeding plumages (Brooke, 1998). The presence of stripes in other social taxa, in particular for the contrasting bands on the bodies of some social mammals, such as gazelles (Caro and Stankowich 2010), and stripes in different species of fishes (Denton and Rowe 1998, Seehausen et al. 1999, Kelley et al. 2013), has similarly been debated but their function remains unclear.

**Fig. 1.**
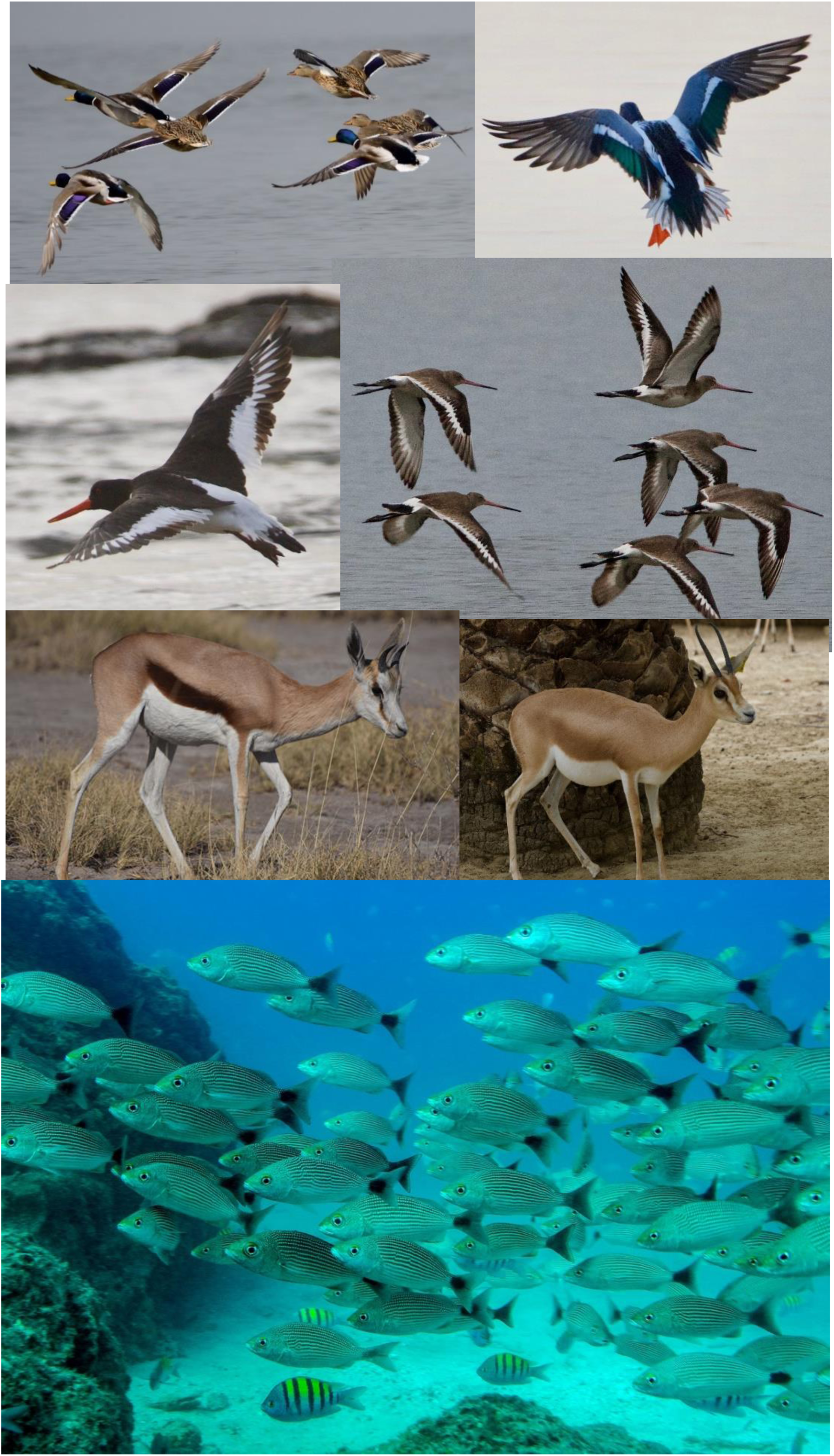
Species of birds, mammals and fishes with examples of contrasting stripes. From left to right, and top to bottom: Mallard *Anas platyrhynchos*, including both males and females; northern shoveler *Spatula clypeata*; common oystercatcher *Haematopus ostralegus*; black-tailed godwit *Limosa limosa*; springbok *Antidorcas marsupialis*; dorcas gazelle *Gazella dorcas*. In the fish picture: with longitudinal stripes, spottail grunts *Haemulon maculicauda*, and with vertical stripes, Panamic sergeant major *Abudefduf troschelii*. Credits: birds and mammals; Juan J. Negro. Fish picture: Christopher Swann.

Others have suggested that such body stripes, when combined in a mass moving group, may produce “dazzle” patterns that could reduce either the likelihood of predators attacking or their attack success rates (Stevens et al. 2008, Hogan et al. 2017, Cuthill 2019). Contrasting stripes or lines have been hypothesized to provide camouflage or to function to startle predators when they are suddenly displayed by fleeing prey (Cuthill 2019, Umbers et al. 2015, Umbers and Mappes 2016). Dazzle patterns may also reduce the ability of a predator to judge the speed and direction of their prey (Hughes et al. 2014). Although there are no empirical data from wild animals, computer simulations using humans as model predators suggest that striped targets are among the most difficult to capture. Nonetheless, simpler patterns, such as all-grey colorations, are equally difficult to catch, and thus evidence that stripes play an adaptive role in dazzle patterning remains inconclusive (Hughes et al. 2014).

Expanding on Brooke’s (1998) original idea for shorebirds, here we separately analyze coloration data on shorebirds, waterfowl, ruminants and fish, while controlling for phylogeny, to assess whether social species are more likely to display stripes on their bodies and investigate their possible function. We posit that social species may use contrasting stripes to aid in coordinated movement or dazzle predators during coordinated movement. According to these hypotheses, we predicted that for any given taxa, contrasting stripes should be more prevalent in group-living species than in solitary ones. Here we test this prediction in species that rely on group cohesion when fleeing from an attacking predator including herding ungulates, flocking birds and schooling fish.

## Material and Methods

### Study species

We selected three different vertebrate taxa: two orders of birds (shorebirds within Charadriiformes, and waterfowl (Anseriformes)), one suborder of mammals (Ruminantia), and fishes in the community of the Eastern Tropical Pacific Ocean with species in the Orders: Anguilliformes, Clupeiformes, Beloniformes, Perciformes, Scombriformes and Istiophoriformes, all of which include species which have contrasting achromatic stripes (i.e., dark and light lines) (Figure 1), as well as other species with no such patterns. We chose to compare the wing stripes in just two bird groups because in these species stripes are only exposed in flight, and thus potentially when the individuals attempt to escape from a predator. Mammals and fish are different in that they do not have foldable limbs. For these groups, we studied the most prominent stripes; longitudinal stripes in mammals, and both longitudinal and vertical in fishes. The taxa under investigation included both solitary and social species that often forage or rest in groups. If approached by a predator, or directly attacked, individuals that detect the predator increase their speed during escape movements, alerting others of the predator’s attack (Ridley et al. 2010, Ridley et al. 2013).

### Charadriiformes

We used the same dataset and species’ ecological attributes as Brooke (1998) who assessed the occurrence of stripes in shorebirds (excluding gulls, auks and allies). Our study included 205 species in this order; although Brooke’s previous study included 210 (we excluded five species because of recent taxonomic changes). Brooke investigated several ecological variables, such as migration, habitat, feeding technique and flocking behavior and found that only flocking behavior was related to the presence of stripes on the wings. For our analysis, we included (a) whether the species was mainly migratory or sedentary, as the marks might help in group forming during migration trips, (b) whether it tended to form flocks or was mainly solitary, and (c) wing chord length as a proxy for overall size. Our response variable was presence/absence of white lines on the wings (1 present, 0: absent). We did not analyze patches on the tails and rumps that were not significantly linked to sociality in Brooke’s (1998) previous study.

### Anseriformes

To capture the difference in stripe expression that we knew *a priori* distinguished the smaller ducks from the larger geese and swans (see Figuerola and Green 2000) we included 148 representative members of all families of waterfowl (this order comprises 180 species in total). We used a binomial response variable indicating the presence or absence of contrasting stripes on the wings following Hegyi et al. (2008). In particular, we unified their scores of 1, 2, and 3 into a single score, i.e. 1 = presence of white patch in the wings, 0 = absence of white patch. For species of geese (not included in Hegyi et al’s (2008) data set), the presence of white wing patch was scored from flying birds displayed in the identification guide of Madge and Burn (1987), and following a similar procedure to Hegyi et al. (2008). For species that are predominantly white, like some swans, elongated dark patches on the wings were scored as presence of stripes.

In contrast to the other taxa included in this study, a majority of the species of Anseriformes are highly social during at least part of their annual cycle. Therefore, for this Order, we could not distinguish between solitary or social species. As an explanatory variable, therefore, we used body mass as a proxy for body size, and a binomial variable indicating if the species is migratory or resident (again to assess whether marks might assist group formation during migrations). We predicted that wing marks would be associated with smaller species (ducks and teals) that are hunted by falcons, whereas these lines should be largely absent in the larger birds (geese and swans). Given the sexual dimorphism in many Anseriformes and the absence of sexual differences in wing patch expression (Hegyi et al. 2008), we selected male body mass to increase variation (female body mass produced similar results – not shown). Body mass data were taken from Figuerola and Green (2000). The migratory status of the species was taken from the distribution maps of the identification guide by Madge and Burn (1987). For partially-migratory species, we classified the species as migratory if the area of breeding range abandoned in winter was larger than the resident breeding area (present all year around).

### Ruminantia

Many species within the Artiodactyla, to which Ruminantia belong, have linear contrasting stripes on their flanks and dorsum (i.e., narrow bands or strips differing in colour from the surface on either side of it). Some species may have other markings of different shapes (i.e., linear or not) in legs, rump and faces. Caro and Stankowich (2010) previously characterized the presence and absence of contrasting marking in different body parts in 198 species of artiodactyls and investigated their function. Here, we built our own dataset to assess whether the longitudinal stripes on the flanks of Ruminantia are related to their degree of sociality. Similar to above, and to remain consistent, we recorded the presence or absence of longitudinal stripes on the bodies of animals as a binary response variable, given the definition of a longitudinal marking by Caro and Stankowich (2010) as a stripe in both flanks contrasting in coloration with the belly below and the dorsum. Social species were defined as the ones forming herds including more than one parent-offspring unit or more than three unrelated individuals. Conversely, solitary species are defined as the ones in which individuals tend to stay alone or just in parent-offspring units. Smaller markings on other body parts (i.e. legs, neck or head) were not considered in the present study, although they may serve a similar function. Body size data were obtained from Smith et al. (2003).

### Fishes

We selected representative species of the main Class of fishes, the Actinopterygii. Among them, we chose species from the fish community of the Eastern Tropical Pacific Ocean, following the guide by Allen and Robertson (1998) and the online-actualized version https://biogeodb.stri.si.edu/sftep/en/pages. Because many tropical species of fish can be colorful, we chose species with several patterns, including species that were both striped and un-striped on the sides of their body. Our searching criteria was as follows: First, and following the phylogenetic order of the book, we identified species with some form of stripe, either parallel to the horizontal axis –i.e, horizontal stripes, or down the vertical axis –i.e., vertical stripes (or bars as in Seehausen et al. (1999)). After this initial selection, we went through our database, and for each species with vertical or horizontal stripes, we randomly identified another species within the same genus, or genus within the same family, that did not have either of those stripes (plain species as in Kelley et al. 2013). In our final database of fishes, most species (90%) belonged to the Perciformes, but with some representatives of additional Orders (Anguilliformes, Clupeiformes, Beloniformes, Scombriformes and Istiophoriformes). Finally, and to align coherently with the other taxa in this study, we only retained primary prey species (defined as species that remain small enough to be eaten by larger fish throughout their lives), and recorded as binary response variables presence/absence of one or more stripes (any color), and their sociality level (i.e., solitary versus schooling).

### Phylogenetic analysis

We tested whether the presence of stripes on the body correlated with the evolution of group living, or other attributes such as body mass or ecological characteristics, using phylogenetic logistic regression (Paradis et al. 2004; Ives & Garland 2010, 2014; Ives et al. 2011; Ho & Ané, 2014; R Core Team 2019). In comparative analyses of trait evolution, both “complete” trees (i.e., those that include unsampled species using polytomy solvers) and “sequenced species only” trees have their advantages and disadvantages (Upham et al. 2019). First, birth-death polytomy resolvers used to infer “complete trees” place species onto trees randomly and independently from the trait’s values, thus potentially breaking down natural patterns of trait phylogenetic structure (Rabosky 2015). Similarly, “sequenced species only” trees, while not suffering from this problem, contain fewer species that are potentially non-randomly sampled, risking excluding their trait values from analyses (Upham et al., 2019). Accordingly, here we followed Upham et al.’s (2019) recommendations and compared analyses run on samples from both complete and sequences only trees (see supplementary material).

Overall, we downloaded phylogenetic subsets from broad phylogenetic syntheses and then summarized those trees by computing single 50% majority-rule consensus trees using SumTrees v 4.4.0 in DendroPy v4.4.0. (Sukumaran & Holder, 2010, 2015) following Rubolini et al. (2015). For the two bird taxa, we downloaded phylogenetic subsets from the BirdTree (Jetz et al. 2012, http://birdtree.org). In particular, we download phylogenetic trees from the Hackett tree distributions (1,000 trees from “All species” + 1,000 trees from “Sequenced species” distributions). For the Ruminantia taxa, we used the species-level trees of extant Mammalia of Upham et al. (2019) (1,000 trees from “birth-death node-dated completed trees” + 1,000 trees from “birth-death node-dated DNA-only” distributions). Lastly, for fish taxa, we downloaded data from the fish tree of life (100 trees from “all-taxon assembled” distributions plus the “DNA-only” time-calibrated tree; Rabosky et al. 2018). The final consensuses “complete trees” were composed of 205 Charadriiformes, 148 Anseriformes, 197 Ruminantia, and 159 fish species, and the final consensuses “sequenced species only” trees were composed by for 144 Charadriiformes, 137 Anseriformes, 182 Ruminantia, and 120 fish species.

To assess whether stripes were associated with group-living behavior, we used phylogenetic logistic regression analyses with the function *phyloglm* (logistic_MPLE method, 1,000 independent bootstrap replicates, and p-values computed using Wald tests) in the phylolm R package v.2.6 (Paradis et al. 2004; Ives & Garland 2010, 2014; Ives et al. 2011; Ho & Ané 2014; R Core Team 2019). We evaluated model fit with binned residual plots –specifically, whether standard error bands from these plots contain 95% of the binned residuals– (*binnedplot* function from the arm v.1.10-1 R package; Gelman & Su 2018) and computing an R^2^ statistic suitable for statistical models with correlated errors, such as the phylogenetic logistic regression models used here (*R2* function from the rr2 R package; Ives 2019). Then, we used the fitted coefficients from the phyloglm models to plot the phylogenetic logistic regression curves using the *plogis* function of the R package “stats.”

## Results

We found significant associations between presence of stripes and the level of sociality in all taxa that we investigated, and these results were consistent across complete and sequenced only trees (see supplementary material). Flocking Charadriiformes (shorebirds) were more likely to have white wing stripes than non-flocking species (phyloglm: R^2^_lik_ = 0.26, *P* = 0.001; Table 1, Fig 2A). Migratory status or body size (wing-chord length), however, were not significantly related to the presence or absence of wing markings (phyloglm: migratory status, *P* = 0.599; body size, *P* = 0.121; Table 1). In Anseriformes (waterfowl), most species of ducks had white wing stripes on dark backgrounds, whereas most of the geese did not (see complete tree in Fig. 2B, and DNA-only tree in Fig. 2BS). In these species, body mass was significantly correlated to the presence or absence of stripes, with smaller species more likely to have stripes than larger ones (phyloglm: R^2^_lik_ = 0.18, *P* = 0.001; Table 1, Fig. 3B). Similar to the Charadriiformes, migratory status was not related to the presence or absence of wing stripes (phyloglm: R^2^_lik_ = 0.18, *P* = 0.346; Table 1). A model excluding the three seemingly outlying data points (i.e., species with body mass > 10,000 g) yielded essentially similar results (phyloglm: R^2^_lik_ = 0.12; body mass, *P* = 0.003; migratory status, *P* = 0.337).

**Table 1.** Summary of phyloglm models.

|  | *Estimate* | *SE* | *95% CI* | *R<sup>2</sup><sub>lik</sub>* | *Z* | *p* |
| --- | --- | --- | --- | --- | --- | --- |
| ***Charadriiformes*** | - | - | - | 0.261 | - | - |
| *α* | 0.015 | - | 0.015, 0.015 | - | - | - |
| *Intercept* | 0.105 | 0.811 | 0.185, 0.845 | - | 0.129 | 0.897 |
| *Sociality* | 0.773 | 0.291 | 0.552, 0.793 | - | 2.659 | 0.008 |
| *Wing chord* | -0.010 | 0.003 | 0.497, 0.500 | - | -1.550 | 0.121 |
| *Migratory status* | -0.025 | 0.047 | 0.470, 0.517 | - | -0.526 | 0.599 |
| ***Anseriformes*** | - | - | - | 0.180 | - | - |
| *α* | 0.016 | - | 0.016, 0.016 | - | - | - |
| *Intercept* | 1.688 | 1.013 | 0.426, 0.975 | - | 1.666 | 0.096 |
| *Body mass* | -0.001 | 0 | 0.500, 0.500 | - | -3.202 | 0.001 |
| *Migratory status* | -0.222 | 0.236 | 0.335, 0.56 | - | -0.943 | 0.346 |
| ***Ruminants*** | - | - | - | 0.273 | - | - |
| *α* | 0.136 | - | 0.031, 0.743 | - | - | - |
| *Intercept* | -1.885 | 0.506 | 0.053, 0.290 | - | -3.728 | 0 |
| *Sociality* | 1.393 | 0.475 | 0.614, 0.911 | - | 2.936 | 0.003 |
| *Body mass* | 0 | 0 | 0.500, 0.500 | - | -1.502 | 0.133 |
| ***Fish (longitudinal)*** | - | - | - | 0.369 | - | - |
| *α* | 0.016 | - | 0.008, 0.133 | - | - | - |
| *Intercept* | -0.548 | 0.342 | 0.228, 0.531 | - | -1.602 | 0.109 |
| *Sociality* | 1.616 | 0.367 | 0.710, 0.912 | - | 4.402 | 0 |
| ***Fish (vertical)*** | - | - | - | 0.162 | - | - |
| *α* | 0.046 | - | 0.024, 0.28 | - | - | - |
| *Intercept* | 0.142 | 0.232 | 0.422, 0.645 | - | 0.612 | 0.541 |
| *Sociality* | -0.9 | 0.346 | 0.171, 0.444 | - | -2.605 | 0.009 |

**Fig 2.**
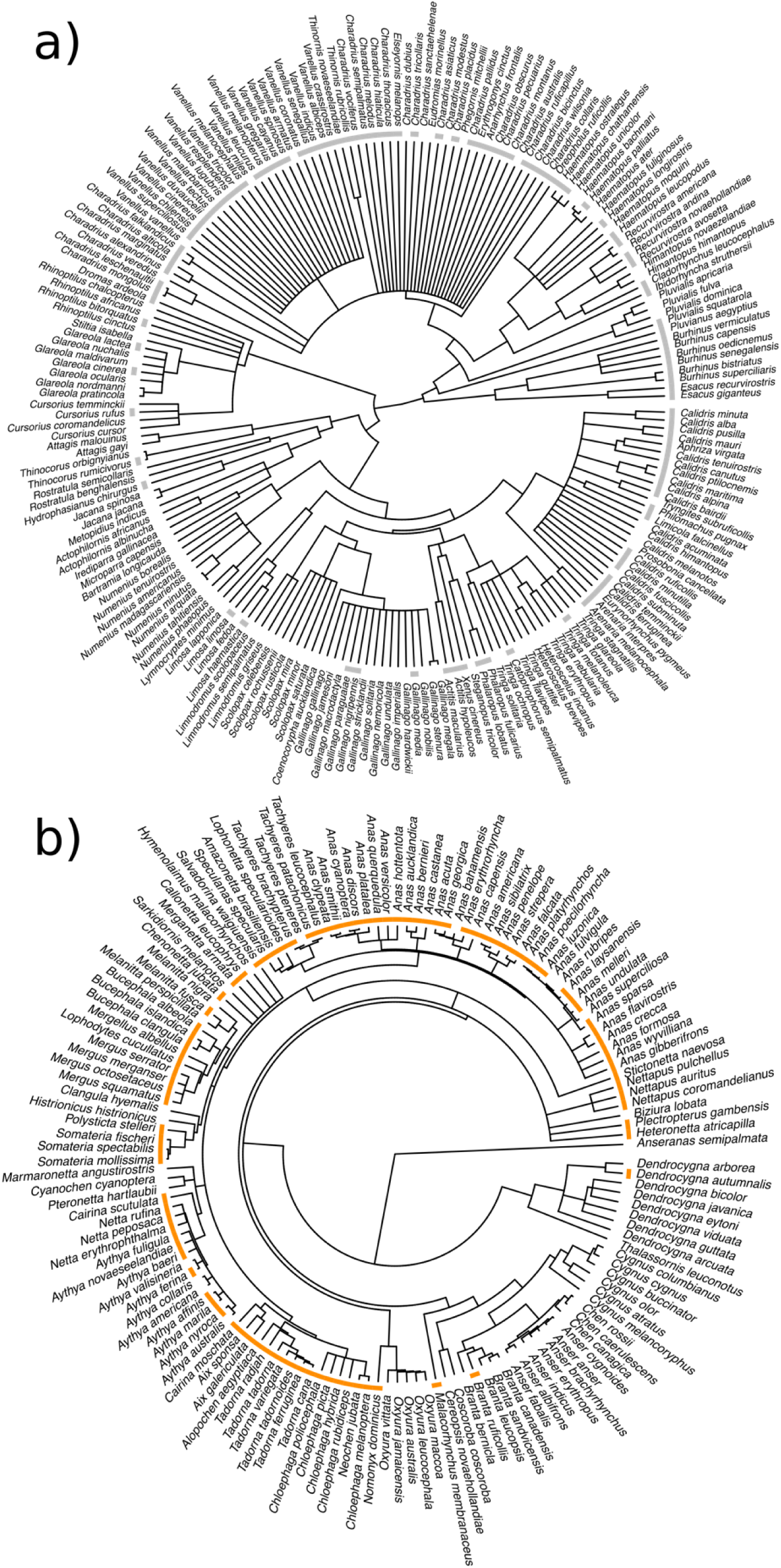
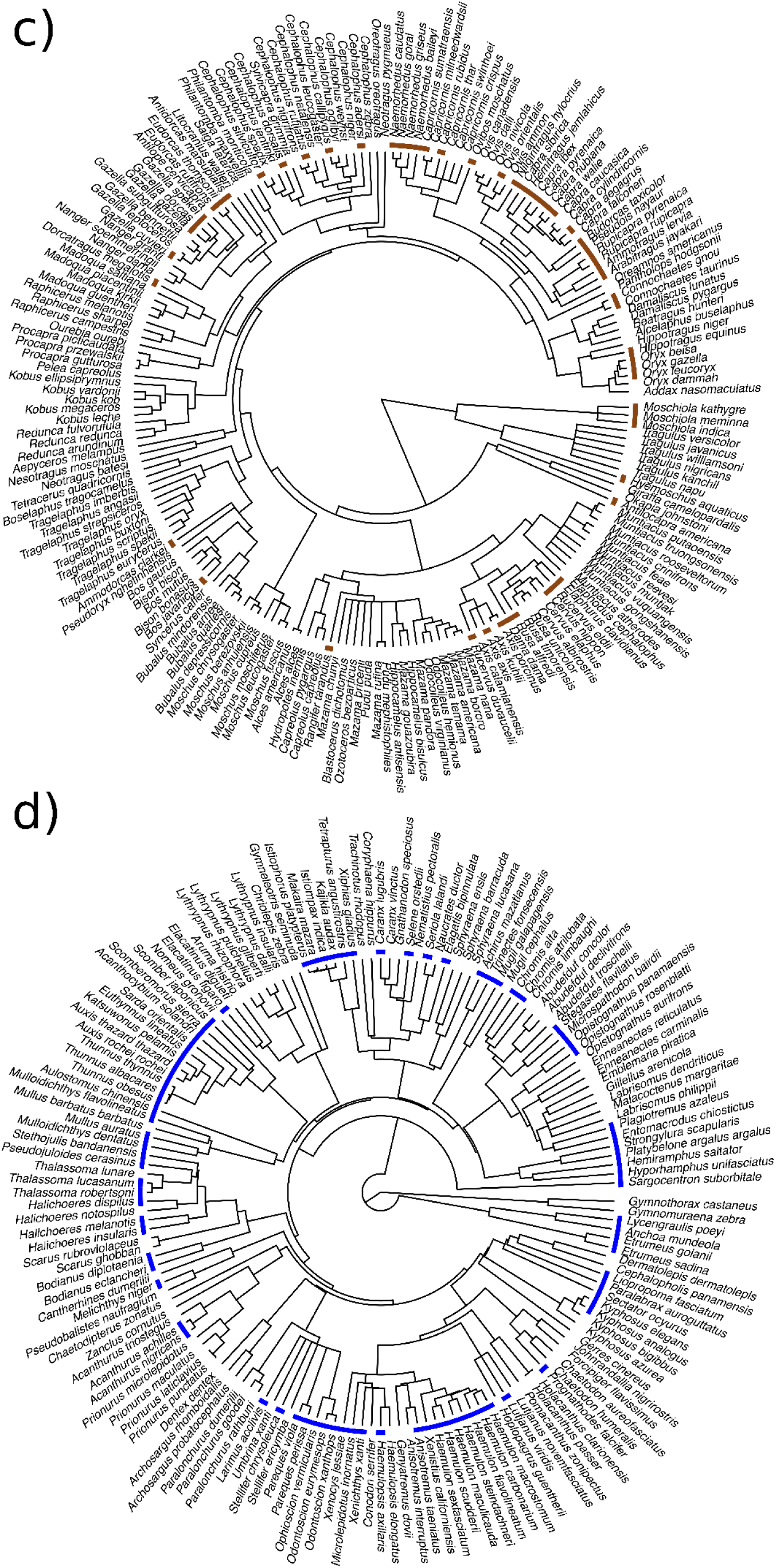
Trait plots depicting the distribution of contrasting stripes (colored traits) across the phylogeny of shorebirds (Charadriiformes, a, gray), Anseriformes (b, orange), Ruminants (c, brown), and Fishes (d, blue).

**Fig 3.**
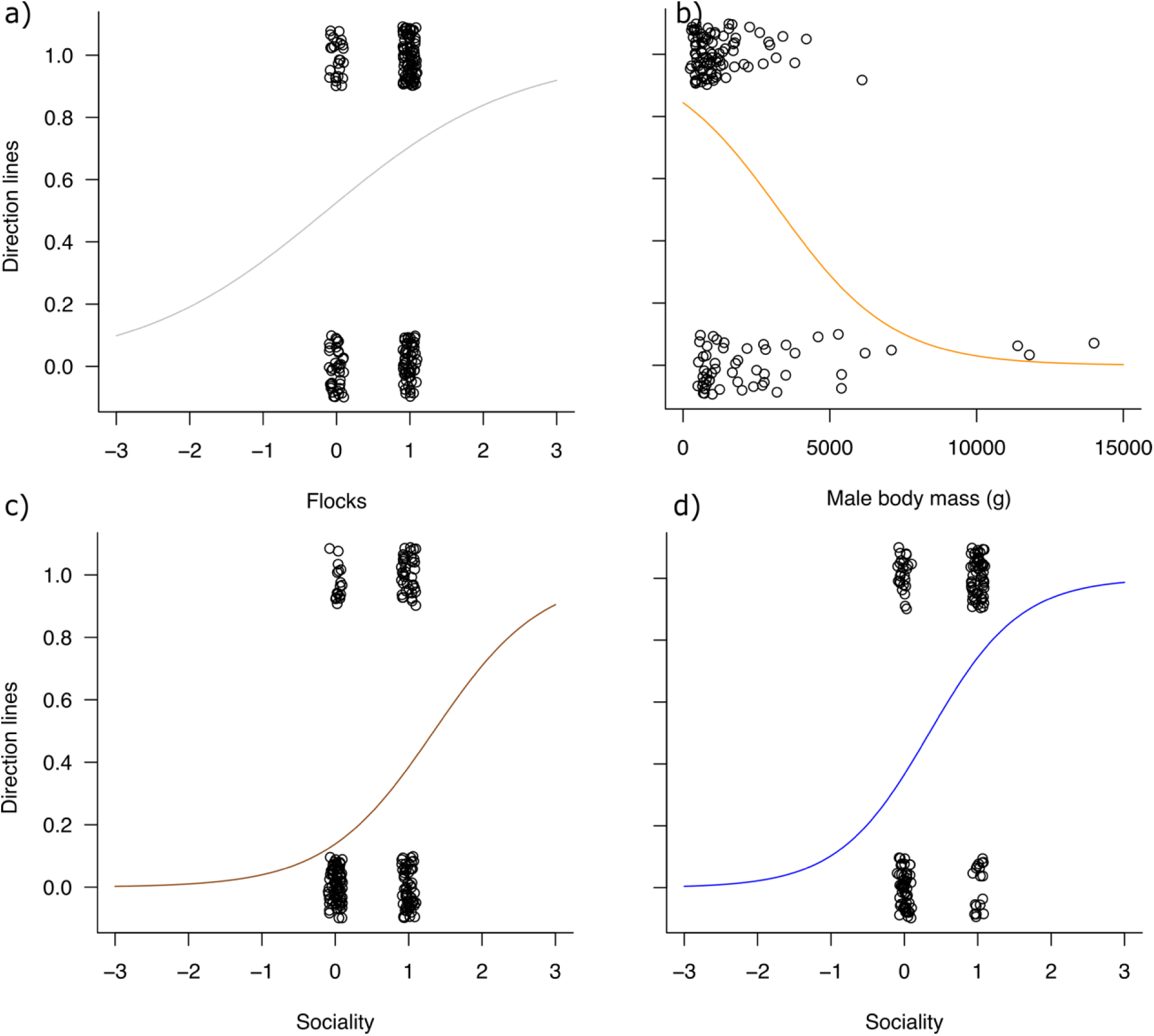
Phylogenetic logistic regression curves for Charadriiformes (a, gray), Anseriformes (b, orange), Ruminants (c, brown), and Fishes (d, blue). Individual points are horizontally and vertically jittered to improve visualization.

Social species of ruminants were also more likely to have stripes on the flanks than non-social species (phyloglm: R^2^_lik_ = 0.27; *P* = 0.003; Table 1, Fig. 3C). Body mass, however, was not related to the presence or absence of stripes (phyloglm: body mass, *P* = 0.133; Table 1).

In fishes, sociality was associated with the presence or absence of horizontal stripes on the body, with social species more likely to have horizontal stripes (phyloglm: R^2^_lik_ = 0.37; *P* < 0.001; Table 1, Fig. 3D). Vertical lines (bars), however, were associated in the opposite direction to social species (i.e., social species were less likely to have vertical stripes; phyloglm: vertical stripes, R^2^_lik_ = 0.16; *P* = 0.009; Table 1).

## Discussion

Stripes on the plumage, fur and scales of prey species were strongly associated with group living across taxa. While different roles for the contrasting marks on animals’ bodies had previously been suggested in independent studies of shorebirds, waterfowl, ruminants and fishes (Brooke 1998, Hegyi et al. 2003, Caro and Stankowich 2010, Kelley et al. 2013, Seehausen et al. 1999), evidence for their importance in social behavior remained inconclusive. In the all-social waterfowl, wing stripes were related to the species’ body mass; a strong predictor of predation risk (Cresswell, 1994, Pomeroy 2006). Given this, and given predation is a strong driver of group living, we suggest these contrasting lines play an important role in social anti-predator defenses of at least three different vertebrate classes; mammals, birds and fishes.

The finding that stripes are associated with social behavior is even more striking considering the visual systems of these taxa are known to vary widely. However, the combination of adjacent achromatic lines or bands – mostly black/dark and white (except in some fishes where structural colors also play a role), suggests that color perception is not necessary for their role. Indeed, detection of patterns and of motion is achieved by achromatic mechanisms, and thus is essentially colorblind (Osorio et al. 1999). These different taxa, therefore, appear to have converged on a similar use of achromatic patterns, as would be expected given signal-detection-theory. Indeed, white bands with no pigment combined with melanized areas may be generated by virtually any vertebrate (McGraw 2006). Hence, these markings may be just as efficiently detected in either di-, tri-, or tetrachromatic species (Bennet and Thery 2007, Bowmaker 1998, Marshall 2000).

While the association between stripes and social behavior appears widespread, these markings’ functional role remains unresolved. Two primary (non-exclusive) functions have been proposed for these stripes. First, they could be used by conspecifics to inform neighbors of rapid movement and aid in coordinated movement (Brooke 1998), or second, to reduce predation success through flash, dazzle, or confusion effects (Stevens et al. 2008, Cuthill 2019). In many cases of birds (including the Charadriiformes and Anseriformes studied here) wing markings are often only exposed when a bird takes flight, but remain hidden when the bird folds its wings, with cryptic dorsal plumages intended for background matching (Cuthill 2019). This sudden display of contrasting lines has led many authors to consider them primarily as flash or dazzle marks destined to confuse an approaching predator that might misjudge the speed or direction of the prey’s escape (Brooke 1998). While such an effect could also be achieved by a single individual exposing its wings during its escape flight, the flashes from multiple individuals would likely produce a stronger stimulus, and could hence be used by social animals to reduce per capita predation risk. Indeed, because predators prioritize distance estimation, their perception of horizontal contours may be disrupted by such movements, viewing such collectively moving prey as blurred images (Banks et al. 2015). While such marking may confuse predators from long-range, at short-range on the other hand, these markings may indicate to other group members the presence of a predator, and hence achieve more coordinated take-offs (Brooke 1998). Of course, the presence of stripes is not restricted to social species. Some solitary species have stripes, but in these cases, these patterns appear to function as aposematic signals, as in skunks (Stankowich et al. 2011), or deimatic signals that startle predators (Umbers and Mappe 2016). However, our analysis suggests that, all things being equal, social species are more likely to have stripes than non-social species.

In the Ruminantia, longitudinal (horizontal) stripes were more likely to be associated with species that formed groups, irrespective of their body size, and horizontal lines in fishes were associated with shoaling. The roles of lines in both these taxa has previously been debated. In cichlid fishes, open-water species that engage in shoaling behavior tend to have longitudinal stripes (Seehausen et al. 1999), but vertical or horizontal stripes in butterflyfishes are not associated with social behavior, while diagonal stripes are negatively correlated with fish’s shoaling tendencies (Kelley et al. 2013). Our work represents definitive evidence that across a wide phylogenetic tree, horizontal stripes indeed appear important in social behavior. In contrast to the wing markings of birds that are only exposed when birds fly, longitudinal lines of mammals and fish are permanently visible. What roles do these markings play, therefore, in social behavior? Caro and Stankowich (2010) suggested several functions for the stripes in Ruminantia, including pursuit deterrence, intraspecific signaling and crypsis. In fishes, some authors have suggested that stripes may disrupt the body shape making it unrecognizable for predators (Armbruster and Page, 1996) or that stripes may make it difficult for a predator to focus on a specific individual target (the confusion effect, McRobert and Brander 1998). Once again, the association of grouping with these stripes suggests that both coordination and confusion effects may play an important role for these species. Indeed, Denton and Rowe (1998) suggested that stripes in the mackerel *Scomber scombrus* help to coordinate shoaling behavior because the way stripes are perceived changes with body orientation (although they referred to the vertical zebra stripes of the upper part of the fish body). Further studies on the functional role of horizontal and vertical stripes are clearly warranted.

Prerequisites for both confusion or coordination hypotheses are that species should be diurnal, and live in relatively open habitat in order to make the signals visible. These assumptions appear to be supported in the Anseriformes and Charadriiformes that often live on open water, marshlands and coastlines, and in the species of ruminants studied which tend to graze on open grasslands (Caro and Stankowich 2010). Indeed, even though many mammals are nocturnal, Stoner et al. (2003) noted that artiodactyls with contrasting lines on their flanks were diurnal and lived in open habitats. The fishes in our sample similarly live in illuminated shallow waters.

While our result confirms many social species have stripes and contrasting marks, there are species which form large moving groups in open environments without any conspicuous contrasting markings. Indeed, the European starling *Sturnus vulgaris*, that does not display visually contrasting patterns in the plumage (Svensson et al. 1999), yet displays some of the most coordinated and collective behaviour in the animal kingdom (Ballerini et al. 2008a, 2008b, Cavagna et al. 2008a, 2008b, 2010). Clearly, stripes are not a prerequisite for coordinated and cohesive movement. Stripes may have other functions, for instance, melanin-based plumage patches have been suggested to signal dominance in social bird species. Nonetheless, these black patches are located in frontal areas such as the head and breast, and are thus assessed in close encounters among conspecifics (Senar 2006).

Using a comparative method where we accounted for phylogenetic signal, we found that contrasting and achromatic stripes are more likely to be present across a broad taxonomic range of social species that are often predated upon. It appears that these stripes may aid in group coordination, or act to increase the confusion effect, both of which may reduce per capita predation risk. These achromatic patterns even suggest that communication signals for alerting and recruiting others may even work within mixed-species groups that are so prevalent among social birds –including ducks and waders-, mammals –including gazelles- and fish (Goodale et al. 2017). Our work represents a rare example of aligned phenotypic characteristics across a wide range of taxonomic diverse animals.

## Acknowledgements

We thank Enrique Figueroa-Luque for his help in building the dataset for mammals, and Christopher Swann for the fish picture. AR was supported by Juan de la Cierva programme, Spanish Ministry of Economy, Industry and Competitiveness (IJCI-2015-23913). JEH-R was supported by the Whitten Lectureship in Marine Biology, and a Swedish Research Council grant number 2018-04076.

