## Supplementary material for "Contrasting stripes are a widespread feature of group living in birds, mammals and fishes": see supplementary material (from another version of this preprint: version 1, 2020-04-20)

Table S1. Summary of phyloglm models using "sequenced species only" trees

|  | **Estimate** | **SE** | **95% CI** | **R^2^_lik_** | **Z** | **p** |
| --- | --- | --- | --- | --- | --- | --- |
| **Charadriiforms** | - | - | - | 0.201 | - | - |
| α | 0.015 | - | 0.015, 0.015 | - | - | - |
| Intercept | 0.688 | 0.952 | 0.235, 0.928 | - | 0.723 | 0.47 |
| Flocks | 0.51 | 0.338 | 0.462, 0.764 | - | 1.509 | 0.131 |
| Wing chord | -0.007 | 0.004 | 0.497, 0.5 | - | -1.876 | 0.061 |
| Migratory status | -0.048 | 0.049 | 0.464, 0.512 | - | -0.981 | 0.326 |
| **Anseriformes** | - | - | - | 0.269 | - | - |
| α | 0.016 | - | 0.016, 0.016 | - | - | - |
| Intercept | 1.49 | 1.038 | 0.367, 0.971 | - | 1.435 | 0.151 |
| Body mass | -0.001 | 0 | 0.5, 0.5 | - | -3.32 | 0.001 |
| Migratory status | -0.102 | 0.263 | 0.35, 0.602 | - | -0.386 | 0.7 |
| **Ruminants** | - | - | - | 0.269 | - | - |
| α | 0.146 | - | 0.025, 0.788 | - | - | - |
| Intercept | -1.885 | 0.506 | 0.053, 0.29 | - | -3.728 | 0 |
| Sociality | 1.393 | 0.475 | 0.614, 0.911 | - | 2.936 | 0.003 |
| Body mass | 0 | 0 | 0.5, 0.5 | - | -1.502 | 0.133 |
| **Fish (longitudinal)** | - | - | - | 0.375 | - | - |
| α | 0.018 | - | 0.008, 0.274 | - | - | - |
| Intercept | -0.766 | 0.365 | 0.185, 0.487 | - | -2.099 | 0.036 |
| Sociality | 1.71 | 0.423 | 0.707, 0.927 | - | 4.041 | 0 |
| **Fish (vertical)** | - | - | - | 0.156 | - | - |
| α | 0.076 | - | 0.026, 0.283 | - | - | - |
| Intercept | 0.2 | 0.272 | 0.417, 0.676 | - | 0.734 | 0.463 |
| Sociality | -0.912 | 0.392 | 0.157, 0.464 | - | -2.33 | 0.02 |

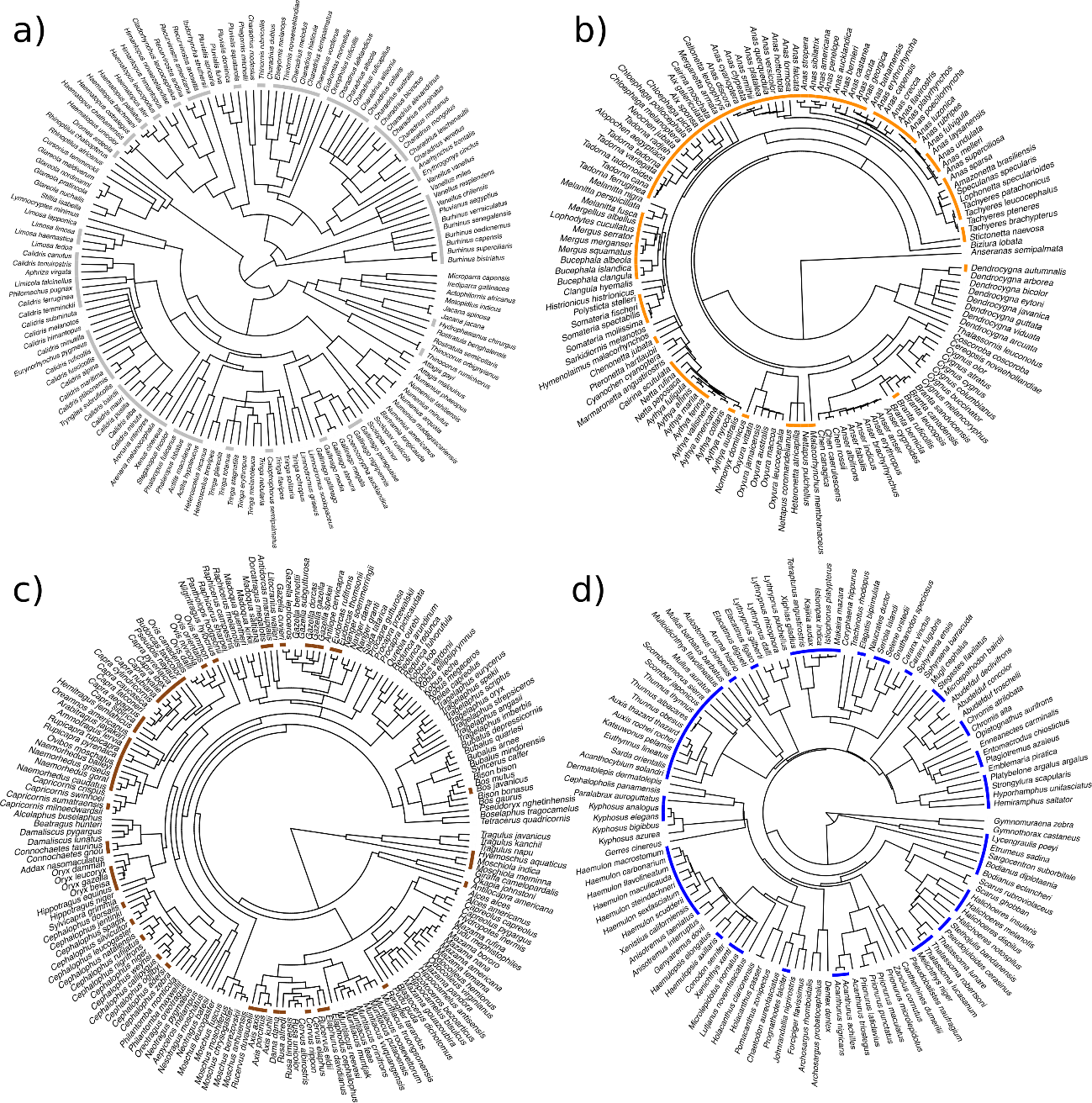

Fig 1S. Trait plots (DNA-only species, see text) depicting the distribution of direction lines (colored traits) across the phylogeny of Charadriiformes (a, gray), Anseriformes (b, orange), Ruminants (c, brown), and Fishes (d, blue).
